# Low oxygen promotes extravillous trophoblast progenitor expansion but restrains maturation

**DOI:** 10.1101/2025.10.29.685387

**Authors:** Gina L. McNeill, Giada Guntri, Ida Calvi, Alastair H. Kyle, Hans-Rudolf Hotz, Victoria M. Leonard, Willie Wu, Matthew J. Shannon, Andrew I. Minchinton, Keegan Korthauer, Margherita Y. Turco, Alexander G. Beristain

**Author notes:** To whom correspondence should be addressed: Alexander G. Beristain, The British Columbia Children’s Hospital Research Institute, The University of British Columbia, Vancouver, British Columbia, Canada. V5Z 4H4., Margherita Y. Turco, Friedrich Miescher Institute for Biomedical Research, Basel, Switzerland. Fabrikstrasse 24, 4056 Basel.

## Abstract

Placental development occurs in a low oxygen environment, yet how oxygen tension instructs differentiation of progenitor cytotrophoblasts (CTB) along the extravillous versus villous pathways remains incompletely understood. In particular, the role of low oxygen in early extravillous trophoblast (EVT) progenitor expansion and subsequent maturation has been difficult to assess due to the lack of models allowing sequential, high-resolution characterization. Using human trophoblast organoids and single-cell transcriptomics, we examined the effects of ambient (21%) and physiological (2–3%, 8%) oxygen on EVT and SCT differentiation. We show that low oxygen promotes the initial expansion of column CTB-like progenitors but restricts terminal EVT maturation. Conversely, we show that ambient oxygen drives the formation of EVT and syncytiotrophoblast. Stabilization of HIF-1 partially recapitulates the EVT maturation block but does not contribute to EVT progenitor expansion, indicating HIF-dependent and-independent contributions. These findings clarify how oxygen shapes EVT lineage progression and define HIF-1’s role in coordinating progenitor expansion versus maturation.

## INTRODUCTION

Villous cytotrophoblasts (CTBs), the progenitor trophoblast population of the placenta, differentiate along two distinct cell pathways: villous and extravillous^1^. Along the villous pathway, CTBs fuse to form a large multinucleated structure called the syncytiotrophoblast (SCT). The SCT functions as the major endocrine engine of the placenta and facilitates nutrient and gas exchange between fetal and maternal circulations^2,3^. The extravillous pathway initiates at placental (villi)-uterine attachment points, where progenitor CTB establish multi-layered proliferative structures called anchoring columns^1,4^. Here, trophoblasts undergo an EMT-like differentiation that culminates in the formation of invasive extravillous trophoblasts (EVT) that detach from anchoring columns and infiltrate uterine arteries, glands, and stroma^5^. EVT, like the multinucleated SCT, are essential for pregnancy success, where EVT participate in uterine artery remodeling and modulate the activity of maternal immune cells to tolerate fetal antigen^6^. Impairments in either the villous or extravillous pathways can lead to placental dysfunction that likely contributes to the development of severe pregnancy disorders like fetal growth restriction^7^ and preeclampsia^8^.

Trophoblasts residing within the proximal portion of the anchoring column, termed column cytotrophoblast (cCTB), are highly proliferative and are described as a source of trophoblast progenitors^9^. Studies using single-cell transcriptomics to characterize placental trophoblasts have identified cCTBs specific to the proximal column as an integrin α2 (*ITGA2*)-expressing population that gives rise to human leukocyte antigen-G (HLA-G)-expressing cCTB representing a more mature phenotype^10^. However, the transcriptional regulators important in initiating the transition of proximal column cCTBs into more mature cCTB has not been clearly defined. While NOTCH1 is a marker of proximal/mid column cCTB and functions to suppress pathways central to CTB self-renewal and cell-cell fusion (while also activating transcriptional programs aligned with EVT development)^11^, its role in primitive EVT progenitor (i.e., integrin α2-expressing cCTB) maintenance has not been examined. Similarly, the HIPPO pathway-associated transcription factors TEAD1 and TEAD3 and their co-factor TAZ induce hTSCs and trophoblast organoids to differentiate into EVT^12,13^. However, like NOTCH1, the developmental timing of their activity in terms of EVT progenitor expansion and maintenance has not specifically been tested. The oxygen-sensing transcription factor hypoxia inducible factor (HIF)1α ^14^ and EPAS1/HIF2α^15^ have also been localized to anchoring trophoblast column, indicating that oxygen-sensing pathways may also in part control early cCTB/EVT progenitor development. In 2-D primary trophoblast cultures, low oxygen promoted the differentiation of CTB to HLA-G-expressing EVT-like cells, and intact HIF-1 was required for this process^16^. Additionally, knockdown of EPAS1 was shown to inhibit EVT differentiation in hTSCs^15^. However, the role of HIF-1 in earlier stages of EVT progenitor expansion and later stages of EVT maturation has not been resolved.

Trophoblasts initially reside within a low oxygen environment, where during the first 10–12 weeks of gestation, oxygen tension in the intervillous space is approximately 20 mmHg (2–3% O₂)^17,18^. As trophoblast-directed uterine artery remodeling advances, intervillous oxygen tension gradually rises to 60 mmHg (∼8% O₂) and is maintained at this level for the remainder of pregnancy^19^. The role of oxygen in regulating trophoblast differentiation has been extensively studied, however the effect of low oxygen on EVT differentiation have yielded conflicting results. While some report that low oxygen promotes EVT differentiation^16,20–22^, others have shown that it restricts EVT differentiation^23–25^. These discrepancies likely reflect differences in trophoblast model systems, inconsistencies in the usage of trophoblast cell-type nomenclature, as well as differences in oxygen levels used to model low oxygen tension (e.g., 1-5%)^26^. Consequently, the precise influence of oxygen on EVT differentiation, and in particular progenitor EVT expansion and maintenance, remains incompletely understood.

To address these gaps, we examined the effects of ambient (21%) and physiological (2–3% and 8%) oxygen levels on trophoblast differentiation using advanced organoid systems that faithfully recapitulate both SCT and EVT development, combined with single-cell transcriptomics to enable the detailed characterization of distinct cell states as progenitor CTB differentiate into EVT. We find that oxygen tension shapes both cell composition and gene expression along SCT and EVT lineages. Consistent with previous reports^25,27^, low oxygen restricts SCT differentiation in both SCT^in^ and SCT^out^ organoid models, the latter which better preserves outer SCT/CTB spatial organization^28^. Importantly, we show that low oxygen drives a marked expansion and stabilization of ITGA2^+^ column-like EVT progenitors, but also partially restricts the formation of mature, invasive EVT populations. Stabilizing HIF under ambient (21%) conditions partly reproduces the EVT maturation block observed in low oxygen. However, HIF activity in the absence of low oxygen does not promote progenitor expansion, indicating that low-oxygen–mediated EVT progenitor maintenance occurs independently of HIF. Together, this work elucidates the role of low oxygen in controlling the development of discrete populations of trophoblasts along the EVT pathway. Moreover, these studies uncover a low-oxygen dependent/HIF-independent process in initiating the earliest steps in EVT differentiation.

## RESULTS

### Low oxygen promotes column progenitor expansion while high oxygen drives endpoint differentiation

Human trophoblast organoids (TOrgs) offer a tractable model that accurately recapitulates progenitor CTB differentiation along the villous and extravillous pathways (Fig. 1A). TOrgs derived from either primary trophoblast preparations (Primary-TOrg)^29^ or from human trophoblast stem cell lines (hTSC-TOrg)^9,30,31^ generate discrete populations of trophoblasts that correspond to CD49b/ITGA2^+^ EVT progenitors, NOTCH1^+^/HLA-G^+^/ITGA5^+^ column-like trophoblasts (i.e., cCTB), and mature HLA-G^+^/NOTCH2^+^/ITGA1^+^ EVT (Fig. 1A). Importantly, these cell types arise sequentially along the EVT differentiation trajectory, reflecting the correct developmental order from progenitor to column-like to mature invasive EVT, establishing the TOrg model as an appropriate tool to study human trophoblast development. Leveraging the temporal and cellular fidelity of hTSC-TOrgs together with single-cell transcriptomics, we set out to interrogate how defined oxygen conditions (3%, 8%, and 21%) influence the sequential progression of EVT progenitors to mature invasive cells over seven days of differentiation (Fig. 1B).

**Figure 1.**
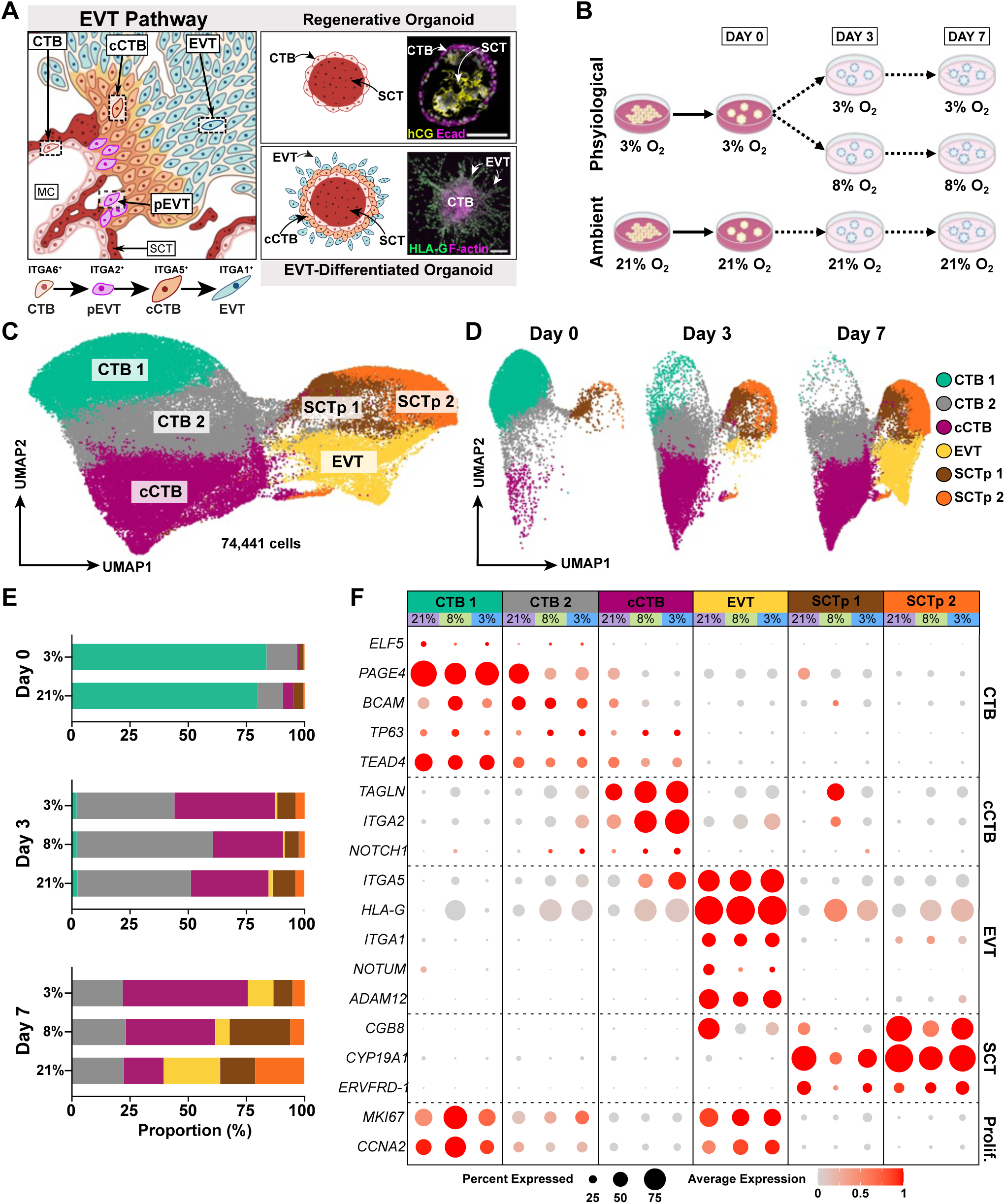
scRNA-seq of hTSC-TOrgs under low and high oxygen. (**A**) (Left) Schematic of a anchoring column structure in the first trimester or pregnancy and the trophoblast subtypes central to the extravillous differentiation pathway. (Right) Schematic and example TOrgs images depicting equivalent trophoblast populations under regenerative or EVT differentiation culture conditions. Organoid are labeled with antibodies targeting hCG, E-cadherin, HLA-G, and filamentous actin (F-actin); nuclei are labelled with DAPI (white). IVS: intervillous space, MC: mesenchymal core; CTB: cytotrophoblast; SCT: syncytiotrophoblast; pEVT: extravillous trophoblast progenitor; cCTB: cell column cytotrophoblast; EVT: extravillous trophoblast. (**B**) Experimental workflow. hTSC-TOrgs were established in 3% or 21% and cultured for 6 days in regenerative media (dark pink) and collected for scRNAseq (Day 0). hTSC-TOrgs were then cultured in EVT media (light pink) for a total of seven days in 3%, 8%, or 21% O_2_. scRNAseq libraries were generated on Day 3 and Day 7 of EVT differentiation in each condition. For all scRNAseq samples, CT29 and CT30 cells were pooled prior to library creation. (**C**) Uniform manifold approximation and projections (UMAP) plot of 74,441 trophoblasts from integrated dataset containing in vitro (21%, 8%, and 3% hTSC-TOrgs) and in vivo dataset. (**D**) UMAP of hTSC-TOrg dataset on each experimental timepoint. (**E**) Proportions of scRNA-seq identified trophoblast clusters in each oxygen condition (21%, 8%, 3%) at each experimental timepoint (Day 0, Day 3, Day 7). (**F**) Dot plot showing the average log expression (0-1) and percent expressed of CTB, cCTB, EVT, and SCT lineage markers, and proliferation (Prolif.) markers in 21%, 8%, and 3% O_2_ in each trophoblast cluster.

hTSC-TOrgs (derived from the CT29 and CT30 hTSC lines^9^) cultured in regenerative (Wnt-potentiating) conditions were first acclimated for 3–4 passages under either 3% or 21% oxygen before initiating EVT differentiation. To confirm that organoids exposed to low oxygen experienced a hypoxic response, we assessed pimonidazole labeling. As expected, organoids maintained in 21% oxygen showed no pimonidazole signal, whereas those cultured in 8% and 3% oxygen displayed a stepwise increase in intensity, indicating that the intended oxygen tensions were achieved (Fig. S1A, S1B). Three differentiation timepoints were collected for single-cell RNA sequencing (scRNA-seq; Days 0, 3, and 7), with Day 0 representing organoids in regenerative media immediately prior to exposure to EVT conditions. To model the physiological transition from low (3%) to moderate (8%) oxygen that occurs near the end of the first trimester, a subset of 3% oxygen–acclimated organoids was switched to 8% oxygen at the onset of differentiation where single cells were collected on Days 3 and 7 (Fig. 1B). By Day 7, organoids in all oxygen conditions exhibited characteristic cellular outgrowth, consistent with successful EVT differentiation (Fig. S1C). Single-cell cDNA libraries were integrated with two published single-cell datasets of first-trimester trophoblasts to aide as a reference for trophoblast subtype identification^30,32^. After preprocessing, filtering, and integration, we identified 61,168 trophoblasts from hTSC-TOrgs and 13,273 from in vivo datasets (Fig. S1D). Clinical characteristics and quality control metrics of all samples are summarized (Table S1).

In the integrated dataset, trophoblasts clustered into six cell states: two CTB-like populations (*ELF5*⁺, *TEAD4*⁺, *BCAM*⁺), a column CTB-like population (*NOTCH1*⁺, *ITGA2*⁺, *TAGLN*⁺), an EVT-like population (*HLA-G*⁺, *ITGA1*⁺), and two SCT precursor (SCTp) populations (*CYP19A1*⁺, *CGB*⁺) (Fig. 1C; Fig S1E; Table S2). Comparing marker gene expression between organoid and *in vivo* trophoblasts, we observed similar patterns across cell states, though with some quantitative differences (Fig. S1E). For example, CTB-associated genes (*ELF5*, *BCAM*, *TP63*) were higher in *in vivo* CTBs (CTB1 and CTB2), whereas cCTB markers (*TAGLN*, *ITGA2*) were more pronounced in hTSC-TOrgs, consistent with prior reports^9^. Despite these expression-level differences, hTSC-TOrgs recapitulate the overall transcriptional landscape and subtype-specific gene patterning of first-trimester trophoblasts, demonstrating the fidelity of the organoid model.

To evaluate how scRNA-seq captures dynamic changes in trophoblast differentiation, we analyzed the proportions of distinct cell states across the 7-day EVT differentiation time course. At Day 0, prior to induction, CTB1 comprised over 75% of all cells, consistent with its role as the most upstream progenitor-like CTB population (Fig. 1D, 1E). By Day 3, the frequency of CTB1 cells declined, while CTB2 and cCTB populations expanded, together representing over 80% of trophoblasts and reflecting the early progression along the EVT lineage (Fig. 1D, 1E). By Day 7, EVT and SCTp2 populations emerged robustly (Fig. 1D, 1E), indicating that the temporal design of our differentiation protocol successfully captured the full spectrum of trophoblast states, from progenitor CTBs to mature EVT and SCT-like cells. Interestingly, the increase in SCTp2 cells highlights the concurrent differentiation of the villous pathway in culture conditions that preferentially drive EVT differentiation. These results establish our scRNA-seq time course as an appropriate platform to assess how discrete oxygen levels impact dynamic EVT differentiation at high cellular resolution.

Building on these baseline dynamics, we next examined how oxygen levels influence the kinetics of specific trophoblast populations. At Day 0, hTSC-TOrgs in regenerative media under 3% and 21% oxygen showed nearly identical frequencies of CTB1 and CTB2 progenitor populations (Fig. 1E). Although column-like cCTBs and SCT precursor (SCTp) populations were minor at this stage, they were slightly more developed in 21% oxygen, suggesting that 21% oxygen modestly promotes early differentiation under regenerative conditions (Fig. 1E). By Day 3, these subtle differences had disappeared, with all three oxygen conditions (3%, 8%, and 21%) exhibiting similar proportions across trophoblast states (Fig. 1E). By Day 7, oxygen-dependent effects became pronounced. Low oxygen (3% and 8%) maintained higher frequencies of column-like cCTBs, whereas ambient oxygen (21%) promoted expansion of mature EVTs (Fig. 1E). Ambient conditions also drove a substantial increase in SCTp2 cells, indicating enhanced villous pathway differentiation. These proportional differences were reflected at the transcriptional level (Fig. 1F): low oxygen (3 and 8%) was associated with elevated expression of progenitor EVT genes (*TAGLN*, *ITGA2*, *NOTCH1*) in cCTB-like cells, while *HLA-G* transcripts were broadly increased under low oxygen in all non-EVT states; these levels, though increased, were still markedly lower than in EVT (Fig. 1F). Conversely, the mature EVT marker, *NOTUM*, was enriched in EVTs under 21% oxygen (Fig. 1F). Collectively, these data indicate that low oxygen expands and/or stabilizes progenitor EVT-like cCTBs, whereas higher oxygen levels favor maturation along both the villous (SCT) and extravillous (EVT) pathways.

### Low oxygen restricts SCT differentiation

Because hTSC-TOrgs cultured in 21% oxygen exhibited a potential enhancement of the villous pathway, we next examined oxygen-dependent differences in SCTp development. Using our hTSC-TOrg scRNA-seq dataset, we compared the expression of a SCT-specific gene signature across SCTp1 and SCTp2 states (collapsed) under 3%, 8%, and 21% oxygen (Fig. 2A). Genes encoding retroviral envelope proteins critical for cell-cell fusion (*ERVW1*, *ERVFRD1*, *ERVV1*, *ERVV2*) were markedly higher in 21% oxygen compared to 3% or 8% conditions. Similarly, genes encoding SCT-associated hormones (*CGA, CGB1, CGB2, CGB5, CGB7, CGB8, INSL4, LEP*), as well as the canonical SCT marker *SDC1*/syndecan1, were also strongly elevated in 21% oxygen (Fig. 2A). Together, these findings indicate that gene programs underlying SCT fusion and hormone production are potentiated under ambient oxygen and, conversely, are suppressed under low oxygen conditions.

**Figure 2.**
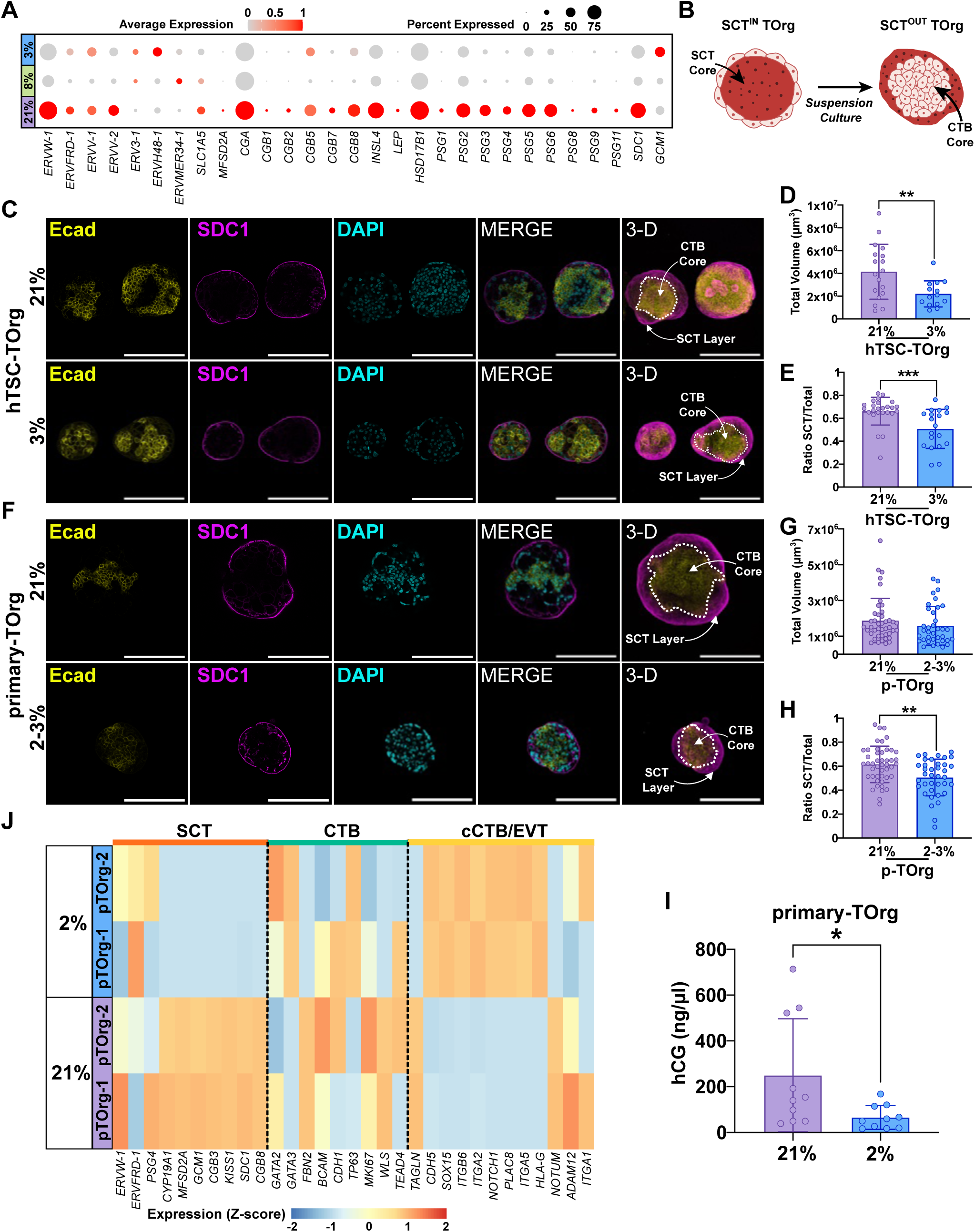
Low oxygen disrupts SCT-associated gene programs and impairs SCT development. (**A**) Dot plot showing the average log expression and percent expressed of SCT marker genes in SCTp cells (SCTp1 and SCTp2, combined) in 21%, 8%, 3% O_2_. (**B**) Schematic depicting the process, including suspension culture, required for converting SCT^IN^ TOrgs into SCT^OUT^ TOrgs. (**C**) Representative confocal microscopy images of SCT^OUT^ hTSC-TOrgs (CT29) cultured in 21% or 3% O_2_ after three days in suspension culture. Organoids are labelled with antibodies targeting the CTB marker E-Cadherin (Ecad), the SCT marker syndecan-1 (SDC1), and nuclear marker DAPI. Dotted lines denoted the CTB core. Scale bars, 200µm. (**D**) Bar graph with individual datapoints showing total organoid volume of SCT^OUT^ hTSC-TOrgs (CT29) cultured in 21% and 3% O_2_. Data plotted as mean values with standard deviation error. Statistical analyses were performed using an unpaired t-test; differences significant at P<0.05. **=P<0.01. (**E**) Bar graph showing the ratio of SCT to total organoid volume of SCT^OUT^ hTSC-TOrgs (CT29) cultured in 21% and 3% O_2_. Data plotted as mean values with standard deviation error. Statistical analyses were performed using a Mann-Whitney test; differences significant at P<0.05. ***= P<0.001. (**F**) Representative confocal microscopy images of SCT^OUT^ Primary-TOrgs cultured in 21% and 2-3% O_2_ after three days in suspension culture. Organoids are labelled with antibodies targeting the CTB marker E-Cadherin (Ecad), the SCT marker syndecan-1 (SDC1), and nuclear marker DAPI. Dotted lines denoted the CTB core. Scale bars, 200µm. (**G**) Bar plot showing total organoid volume of SCT^OUT^ Primary-TOrgs cultured in 21% and 2-3% O_2_. Data plotted as mean values with standard deviation error. Statistical analyses were performed using a Mann-Whitney test; differences significant at P<0.05. (**H**) Bar plot showing the ratio of SCT to total organoid volume of SCT^OUT^ Primary-TOrgs cultured in 21% and 2-3% O_2_. Data plotted as mean values with standard deviation error. Statistical analyses were performed using a Mann-Whitney test; differences significant at P<0.05. **= P<0.01. (**I**) Bar plot showing hCG concentration of conditioned media in SCT^OUT^ Primary-Orgs cultured in 21% and 2% O_2_ over 3 days. Data plotted as mean values with standard deviation error. Statistical analyses were performed using a Mann-Whitney test; differences significant at P<0.05. *=P<0.05. (**J**) Heatmap showing the expression level (Z-score) of SCT, CTB, and cCTB/EVT lineage markers in two SCT^OUT^ Primary-TOrgs (pTOrg-1 and pTOrg-2) cultured in 21% and 2% O_2_.

Single-cell RNA-seq cannot capture true multinucleated syncytial structures, limiting its ability to fully assess SCT development. To address this limitation, we leveraged recently developed apical-out (SCT^out^) Torgs^28^, which form an SCT layer on the outer surface of the organoid mimicking the *in vivo* spatial orientation of CTBs and SCT and therefore providing a more physiological relevant context to evaluate the impacts of oxygen on SCT development (Fig. 2B). In contrast, traditional Matrigel-embedded organoids (SCT^in^) develop their SCT within the organoid core where oxygen tension is lower even in ambient conditions^33^, limiting their use for examining the impacts of oxygen on the villous pathway. Thus, to directly assess oxygen effects, SCT^out^ hTSC-TOrgs were cultured under low (3%) and ambient (21%) oxygen in suspension for 3 days. At endpoint, wholemount organoids were probed with antibodies targeting E-cadherin and Syndecan-1/*SDC1* to aide in the visualization and quantification of the CTB core (E-cadherin^+^) and outer SCT layer (Syndecan-1^+^) (Fig. 2C). Organoids maintained in 3% oxygen exhibited a pronounced decrease in total organoid volume (Fig. 2D). Further, SCT^out^ hTSC-TOrgs cultured in 3% oxygen also showed a decrease in SCT volume/total organoid volume ratio (Fig. 2E) indicating that low oxygen impairs SCT formation.

To validate that these oxygen-dependent effects on the SCT are not unique to hTSC-derived organoids, we examined primary-TOrgs (also cultured as SCT^out^) under low (2%) and ambient (21%) oxygen (Fig. 2F). Although the low oxygen condition in these primary organoids was 2% (slightly lower than the 3% used for hTSC-TOrgs), both conditions fall within the physiological range of early first-trimester placental oxygen tension^21,34^. While total organoid volume was not affected by oxygen condition, similar to SCT^out^ hTSC-TOrgs, low oxygen resulted in significantly less SCT relative to total organoid volume, confirming that low oxygen restricts SCT development in this model (Fig. 2G, 2H). Consistent with these findings, hCG secretion was also decreased under low oxygen, indicating functional impairment of the SCT (Fig. 2I).

To gain insight into the transcriptional basis of these effects, we performed bulk RNA-seq on SCT^out^ primary-TOrgs cultured in 2% or 21% oxygen (n=2 donors). Principal Component Analysis showed that organoids clustered strongly by oxygen condition (Fig. 2SA), demonstrating a major impact of oxygen on global gene expression. Focusing on SCT-associated genes, low oxygen reduced expression of key regulators including *GCM1*, essential for initiating SCT development (Fig. 2J)^25^. In line with our scRNA-seq data generated in SCT^in^ hTSC-TOrgs, low oxygen also led to the enrichment of progenitor EVT-associated genes (*SOX15, ITGA2, NOTCH1*), whereas genes linked to mature EVT (*NOTUM, ADAM12*) were elevated under ambient oxygen (Fig. 2J). These results suggest that low oxygen impairs SCT formation by dysregulating early transcriptional pathways while potentially biasing progenitor CTBs toward early EVT-like states.

### Low oxygen potentiates the formation of progenitor EVT but blunts maturation

To address our initial findings showing that low oxygen (3% and 8%) leads to the possible expansion of column-like cCTBs and restricts the emergence of mature EVT-like cells, we leveraged our hTSC-TOrg single-cell dataset to examine EVT differentiation across the full spectrum of cell states contributing to the EVT lineage. To do this, trophoblasts initiating from the earliest CTB-like population (CTB1) and extending to the mature EVT endpoint (i.e., CTB2, cCTB, EVT) were subset and subjected to Monocle3^35^ pseudotime trajectory analyses (Fig. 3A). As expected, this identified the CTB1 state as the differentiation origin and the EVT state as the endpoint, with CTB2 and cCTB states as sequential intermediates along this trajectory (Fig. 3A).

**Figure 3.**
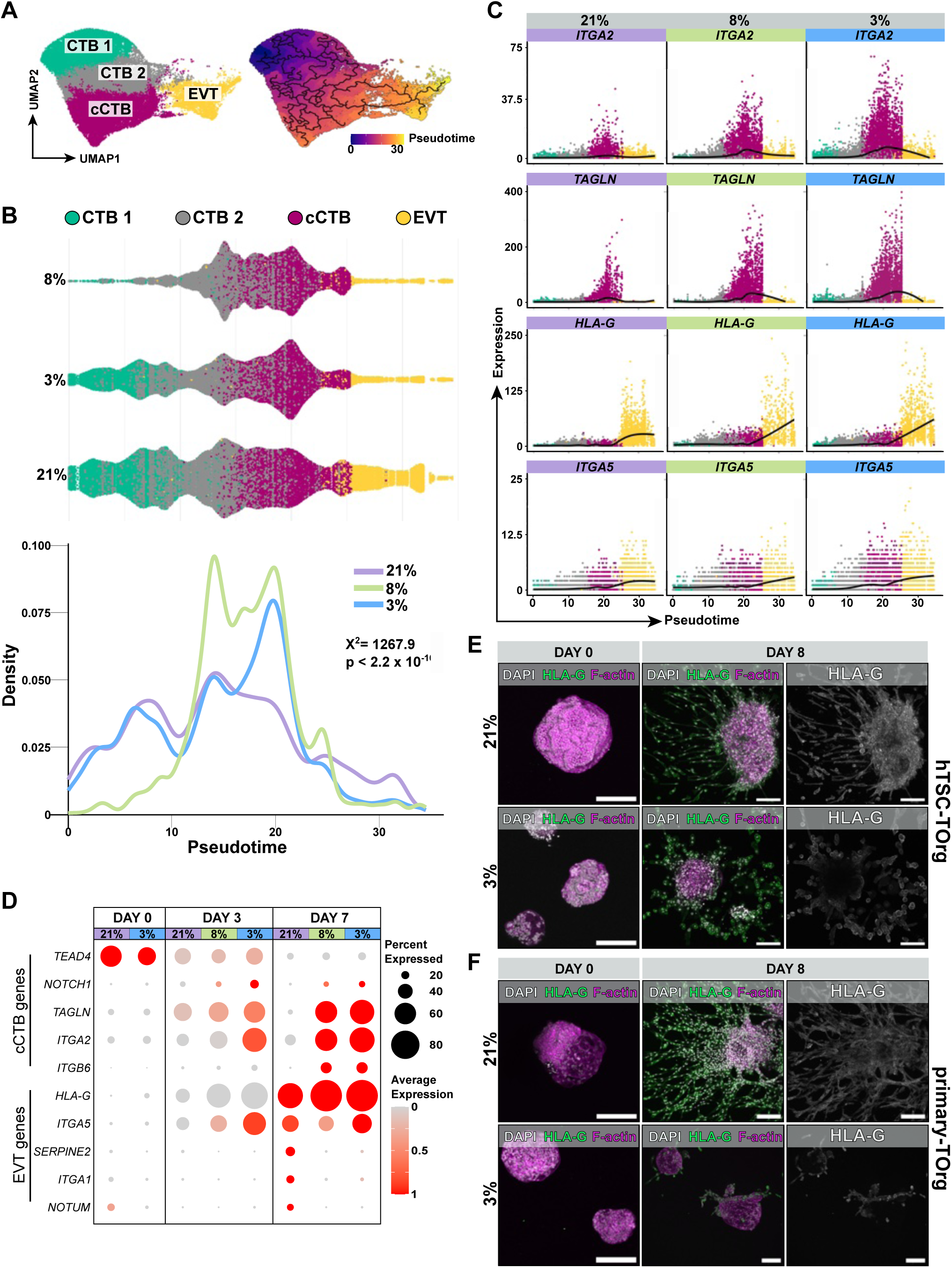
Low oxygen drives progenitor EVT expansion but restricts EVT maturation. (**A**) UMAP of EVT lineage subset (CTB 1, CTB 2, cCTB, EVT) in hTSC-TOrg dataset (21%, 8%, 3%) and UMAP of EVT lineage subset overlain with constructed Monocle3 lineage trajectories (black lines) colour coded by pseudotime values (0-35). (**B**) Swarm and density plots showing cell densities of CTB 1, CTB 2, cCTB, and EVT along the Monocle3 predicted EVT trajectory in 21%, 8%, and 3% O_2_. Dots are colour coded by cell type, and lines are colour coded by condition. Statistical analyses between pseudotime distributions were performed using a Kruskal-Wallis test; differences significant at P<0.05. (**C**) Scatter plot showing gene expression levels (counts) of cCTB (*ITGA2*, *TAGLN*) and early EVT (*HLA-G*, *ITGA5*) transcripts across predicted pseudotime (0-35) in 21%, 8%, and 3% O_2_. Cells are colour-coded by cell state. (D) Dot plot showing the average log expression (0-1) and percent expressed of CTB (*TEAD4*), cCTB (*NOTCH1, TAGLN, ITGA2, ITGB6*) early EVT (*HLA-G, ITGA5*), and late EVT (*SERPINE2, ITGA1, NOTUM*) marker genes in 21%, 8%, 3% O_2_ across each experimental timepoint (Day 0, Day 3, Day 7). Representative confocal microscopy images of (**E**) hTSC-TOrgs (CT29) or (**F**) primary-TOrgs cultured in 21% and 3% O_2_ on Day 0 and Day 8 of an EVT outgrowth assay. Organoids are labelled with antibodies targeting F-actin, HLA-G and DAPI. Scale bars, 200µm.

Examination of cell distributions across pseudotime revealed distinct oxygen level-related differentiation dynamics (Fig. 3B). While the density of progenitor CTB-like cells (CTB1 & 2) was similar between 3% and 21% conditions during early pseudotime values, a significant increase in cCTB (i.e., a mixture of EVT progenitors and column-like cells) is clearly observed in low oxygen at midpoint pseudotime (Fig. 3B). Because trophoblasts specific to the 8% oxygen condition did not factor in until Day 3 of EVT differentiation, they lacked the origin-associated CTB1 state and can only be compared to 3% and 21% mid-pseudotime and later. To this end, 8% oxygen showed marked, but transient increase in CTB2 compared to 3% and 21% conditions (Fig. 3B). However, following this initial spike, cell density dynamics along the trajectory in 8% aligned closely with 3% oxygen, suggesting mid-late EVT differentiation is not significantly different between these conditions (Fig. 3B). Notably, cell density values near pseudotime endpoint (contributed by cells of the EVT state) were greater in 21% oxygen compared with 3% and 8% conditions (Fig. 3B). In addition to overall EVT differentiation kinetics, we investigated the expression of individual genes across pseudotime in each of our conditions (Fig. 3C). Genes associated with a cCTB phenotype (*ITGA2* and *TAGLN*) and early stages of EVT differentiation (*HLA-G*, *ITGA5*), were expressed earlier in pseudotime in 3% and 8% compared to 21% O2, where the initial induction of these genes aligned to cells of the CTB2 and cCTB states (Fig. 3C). Additionally, there was overall higher expression of these markers in 3% and 8% compared to 21% at equivalent pseudotime values (Fig. 3C). 8% represented an intermediate condition, where although expression was earlier and higher compared to 21%, it was lower than cells in 3% oxygen (Fig. 3C). In summary, single-cell pseudotime trajectory modelling showed that low oxygen (3% and 8%) accelerated the expression of genes associated with EVT progenitor and immature EVT development, led to an expansion of column-like cCTB trophoblasts, and restricted the progression and/or formation of mature EVT.

Because pseudotime is only a computationally inferred measure of differentiation, we next examined the expression of genes associated with cCTB and mature EVT states over our three EVT differentiation timepoints (Day 0, Day 3, and Day 7) (Fig. 3D). The cCTB associated genes *NOTCH1*, *TAGLN*, *ITGA2*, and *ITGB6* in 3% and 8% conditions showed earlier induction at Day 3 of EVT differentiation, and by Day 7 showed sustained levels of expression compared to 21% trophoblasts that exhibited an overall dampening in the levels of these EVT progenitor-associated genes (Fig. 3D). In contrast, expression levels of genes associated with a mature EVT phenotype (i.e., *SERPINE2*, *ITGA1*, *NOTUM*) at Day 7 were markedly higher in 21% oxygen compared to levels in either the 3% or 8% conditions (Fig. 3D). These findings support our single-cell trajectory analyses above that together indicate that low oxygen leads to an earlier and more profound expansion of progenitor EVT but restrains their further differentiation into mature EVT.

Our finding that low oxygen retrains the maturation of EVT suggests that low oxygen may also lead to impaired trophoblast motility. To address this, hTSC-TOrgs were subjected to an EVT outgrowth/migration assay where organoids are cultured in regenerative suspension conditions (SCT^out^) and placed onto a thin Matrigel matrix and cultured for 8 days in EVT differentiation media^28^ in either 3% or 21% oxygen. By Day 8 differentiation endpoint, robust outgrowth of HLA-G^+^ cells was observed in organoids cultured in 21% oxygen, whereas in 3% conditions, outgrowth was dampened (Fig. 3E; Fig. S3A). Notably, exposure to low or high oxygen also affected the cellular morphology of migrating HLA-G^+^ cells; cells in 21% adopted an elongated spindle-shaped phenotype characteristic of the morphology observed during 2D EVT differentiation^31^, whereas cells in 3% oxygen appeared more cuboidal and adherent, forming tightly packed clusters consisting of multiple cells (Fig. 3E). To validate these findings, we subjected primary-TOrgs to the same EVT outgrowth assay. Consistent with our findings in hTSC-derived organoids, primary-TOrgs EVT outgrowth was enhanced in ambient oxygen whereas in 3% oxygen, outgrowth was substantially reduced (Fig. 3F; Fig. S3B). While HLA-G^+^ cells in 21% oxygen adopted an elongated phenotype similar to hTSC-TOrg EVT, we did not observe the same adherent and rounded phenotype in hTSCs (Fig. 3F). Together, these data show that low oxygen culture interferes with the development of invasive EVT-like cells, possibly by slowing down the progression of cCTB-like cells differentiating into mature EVT.

### EVT development activates hypoxia-associated programs

To define gene pathways enriched along the EVT trajectory and their oxygen-specific activity, we applied Single-Cell Pathway Analysis (SCPA)^36^ to CTB1 and EVT states, representing the origin and endpoint of the EVT lineage, respectively. SCPA measures pathway activity at single-cell resolution, including directionality and effect size. CTB1-specific pathways enriched *in vivo* and also observed in 3% and 21% oxygen hTSC-TOrgs included Oxidative Phosphorylation and MYC signaling (V1), while 21% oxygen additionally recapitulated G2M Checkpoint, Myc (V2), and E2F pathways enriched *in vivo*, suggesting that ambient oxygen better preserves progenitor CTB programs (Fig. 4A, 4B; Table S3). Importantly, pathway enrichment in progenitor CTBs was largely independent of oxygen (Fig. 4A).

**Figure 4.**
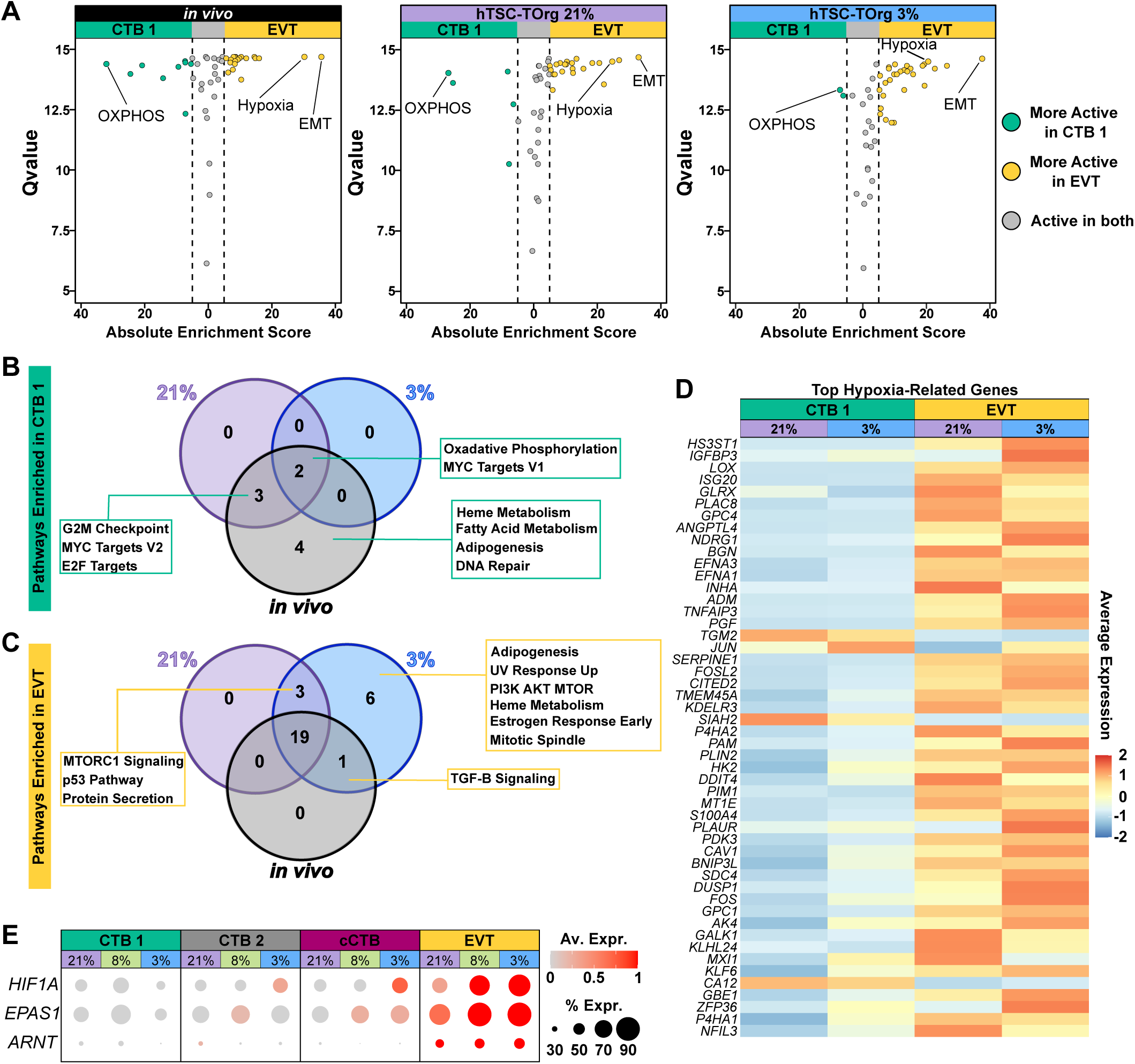
Hypoxia underlies EVT development independent of oxygen condition. (**A**) Single-cell pathway analysis comparing *in vivo*, 21% oxygen, and 3% oxygen CTB 1 and EVT datasets using the Hallmark gene set. Q value measures pathway activity and enrichment score and indicates directionality and effect size of the enrichment. Turquoise dots represent pathways enriched in CTB 1 relative to EVT, and yellow dots represent pathways enriched in EVT relative to CTB 1. Grey dots represent pathways that are active in both CTB 1 and EVT cells. Select pathways are highlighted; oxidative phosphorylation (OXPHOS), Hypoxia, and Epithelial to Mesenchymal Transition (EMT). (**B**) Venn diagram showing pathways that are specific to and shared between in vivo, 21%, and 3% CTB 1 cells. (**C**) Venn diagram showing pathways that are specific to and shared between in vivo, 21%, and 3% EVT cells. (**D**) Heatmap of the top 50 hypoxia-related genes from the DGE analysis of CTB 1 and EVT cells in 21% and 3%. Genes are arranged by absolute fold change. (**E**) Dot plot showing the average log expression (0-1) and percent expressed of the HIF-1 subunits (HIF1A, EPAS1, and ARNT) in CTB 1, CTB 2, cCTB, and EVT cells in the 21%, 8%, and 3% datasets.

In EVT, 19 pathways were consistently enriched across *in vivo* and hTSC-TOrg EVT states from both 3% and 21% oxygen, including EMT, interferon gamma, complement, IL2, glycolysis, and notably, the hypoxia pathway (Fig. 4A, 4C; Table S3). This indicates that hypoxia-related signaling is intrinsic to EVT development. Seven pathways were uniquely enriched in 3% oxygen EVT, including TGF-β (shared with *in vivo* EVT) and six others (adipogenesis, UV response, PI3K-AKT-mTOR, heme metabolism, estrogen response early, mitotic spindle), which may in part contribute to the adhesive, cobblestone-like phenotype observed under low oxygen (Fig. 3F).

Interestingly, hypoxia pathway enrichment was not higher in 3% oxygen EVT relative to 21%, suggesting that activation of hypoxia-related programs is largely oxygen-independent during EVT development. To identify the genes driving this enrichment, we performed differential gene expression on CTB1 and EVT states and ranked the Hallmark “Hypoxia” genes by absolute fold change. EVT in both 3% and 21% oxygen conditions showed a marked induction of most of these hypoxia-related genes compared to the CTB1 state (Fig. 4D). These included genes previously shown to play roles in EVT differentiation (i.e., *LOX*^22^, *PLAC8*^37^, *FOS*^38^, and *ADM*^39^) (Fig. 4D).

Given that many genes in the hypoxia gene set contain HIF-1 binding sites, we next examined HIF subunit expression along the EVT differentiation trajectory. Both HIF1A and EPAS1 increased progressively from CTB1 → CTB2 → cCTB → EVT (Fig. 4E). Within cCTBs, HIF1A was elevated in 3% oxygen, while EPAS1 increased in both 3% and 8% conditions; both remained low in 21% oxygen. As cells matured into EVT, *HIF1A*, *EPAS1*, and *ARNT* were upregulated independent of oxygen, though expression was higher under low oxygen. These findings suggest that HIF-related signaling contributes to both progenitor and mature EVT development, via oxygen-dependent and - independent contexts.

### HIF-1 partially restricts EVT maturation, but does not drive progenitor EVT expansion

Our single-cell analyses revealed that low oxygen (3% and 8%) promotes the expansion of column-like cCTBs while concurrently limiting the emergence of fully mature EVTs (Figs. 1 and 3). Trajectory and gene expression analyses further indicated that hypoxia-associated programs, including HIF-target genes, are robustly activated as cells progress along the EVT lineage, independent of ambient oxygen (Fig. 4). Together, these observations suggest that oxygen tension regulates both the maintenance of immature EVT progenitors and the timing of EVT maturation. Although HIF-1 is known to be required for EVT differentiation^15,16^, its role in specific stages of EVT maturation remains unexplored. We therefore next set out to explore how HIF activity contributes to both the expansion of cCTBs and the restriction of EVT maturation under low oxygen.

To determine which aspects of low-oxygen regulation of EVT differentiation are mediated by HIF signaling, we stabilized HIF1α and EPAS1 (HIF2α) in 21% oxygen using Roxadustat (ROX) — a prolyl hydroxylase inhibitor that prevents HIF1α/2α degradation under normoxic conditions (Fig. 5A). This approach isolates the contribution of HIF activity from other oxygen-sensing mechanisms, allowing us to test whether HIF stabilization is sufficient to drive cCTB expansion or limit EVT maturation independent of low oxygen. hTSC-TOrgs were differentiated in the presence or absence of ROX (50 µM)^40^ for 7 days, with DMSO as a vehicle control and hTSC-TOrgs cultured in 3% oxygen serving as a reference for low-oxygen effects on progenitor expansion and EVT maturation (Fig. 5B). Immunoblotting showed that HIF1α and EPAS1 protein levels were detectable prior to EVT differentiation (Day 0) in 3% oxygen; levels were undetectable in 21% at Day 0 (Fig. 5C). At endpoint EVT differentiation, both HIF1α and EPAS1 levels increased in 3% oxygen, whereas only an EPAS1 lower molecular product (70 kDa) was detected in ambient conditions (Fig. 5C). This latter observation is consistent with a prior report showing that EPAS1 levels increase during EVT differentiation in hTSCs cultured in 21% oxygen^15^. Importantly, treatment with ROX in ambient conditions led to robust stabilization of both HIF1α and EPAS1 (Fig. 5C).

**Figure 5.**
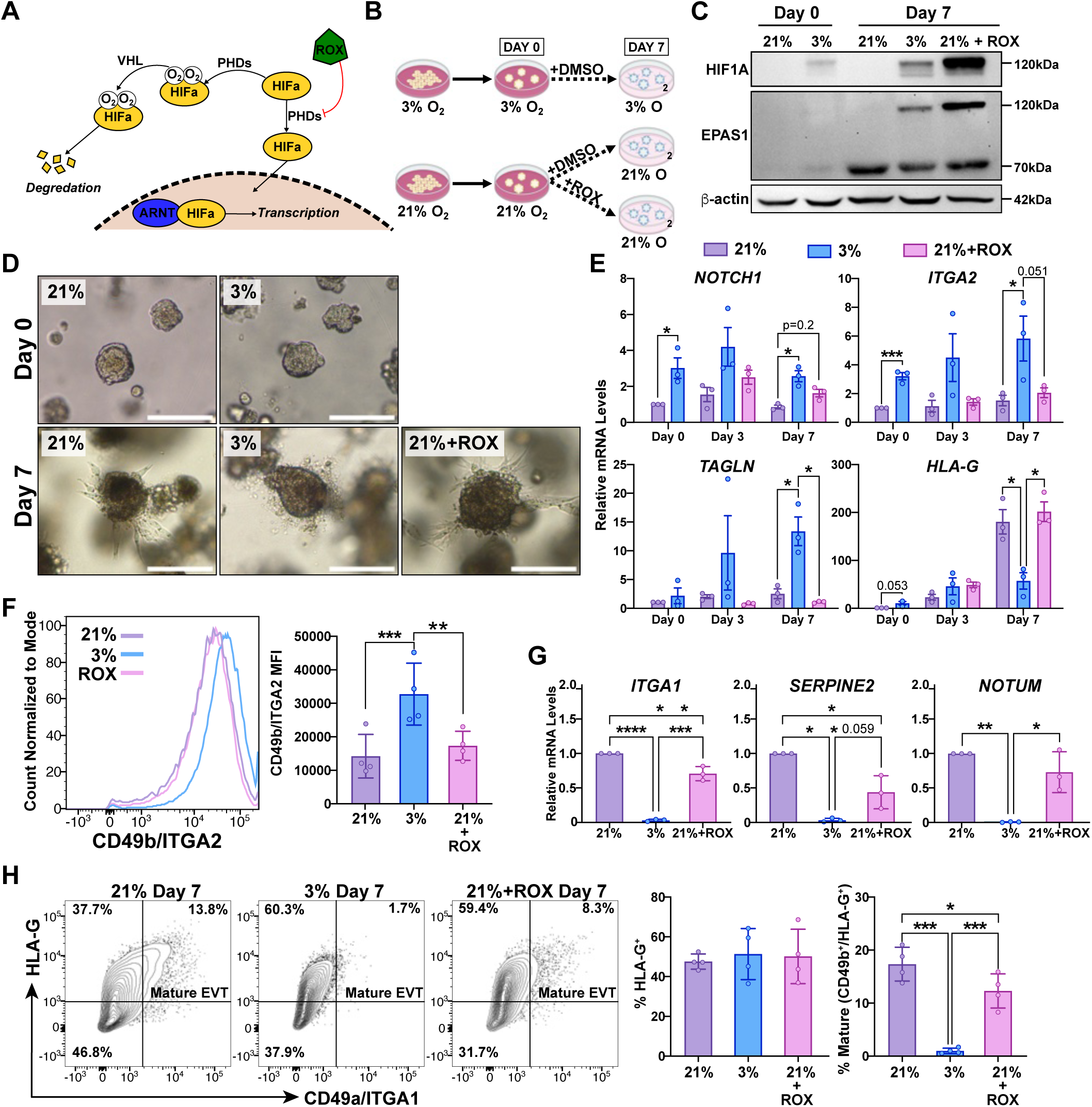
HIF activation alone does not expand progenitor EVTs. (**A**) Schematic of the HIF signalling pathway. In ambient oxygen the HIFα subunits are rapidly degraded, however when oxygen tension is low prolyl hydroxylases (PHDs) cannot hydroxylate HIFα, allowing it to stabilize and bind to ARNT forming an active transcription factor. The pharmaceutical Roxadustat (ROX) stabilizes the HIFα subunits (HIF1α and EPAS1) by blocking the activity of PHDs. (**B**) Experimental workflow. hTSCs (CT29) were established into organoids and cultured for six days in regenerative media in 3% or 21% O_2_. Organoids were then cultured in EVT media for a total of seven days with the addition of either ROX or a vehicle control (DMSO) in 3% or 21% O_2_. (**C**) Immunoblot showing HIF1α and EPAS1 protein levels in hTSC-TOrgs on Day 0 and Day 7 of EVT differentiation in either 3%, 21%, or 21%+ROX conditions. β-Actin levels serve as loading control. Molecular weights (kDa) are indicated. (**D**) Representative brightfield images of hTSC-TOrgs in 21%, 3%, and 21%+ROX on Day 0 and Day 7 of EVT differentiation. Scale bars, 200µm. (E) Relative transcript levels of cCTB (*NOTCH1, ITGA2, TAGLN*), and the EVT lineage (*HLA-G*) in 21%,3%, and 21%+ROX conditions over 7 days of EVT differentiation. Statistical analyses between groups were performed on each timepoint (Day 0, Day 3, and Day7) using either unpaired t-test or ANOVA and two-tailed Tukey post-test; differences significant at P<0.05.*= P<0.05, ***= P≤0.001. (**F**) Representative flow cytometry histogram of ITGA2/CD49b surface expression in 21%, 3%, and 21%+ROX conditions on Day 7 of EVT differentiation (Left). Bar with scatter plot showing the median fluorescent intensity (MFI) of CD49b/ITGA2+ cells in 21%, 3%, and 21%+ROX on Day 7 of EVT differentiation (Right). Data plotted as mean values with standard deviation error. Statistical analyses were performed ANOVA and two-tailed Tukey post-test; differences significant at P<0.05. **= P<0.01,***= P<0.001. (**G**) Relative transcript levels of mature EVT (*ITGA1, SERPINE2, NOTUM*) mRNA in 21%, 3%, and 21%+ROX conditions on Day 7 of EVT differentiation. Statistical analyses between groups were performed ANOVA and two-tailed Tukey post-test; differences significant at P<0.05.*= P<0.05, **= P<0.01,***= P<0.001, ****=P<0.0001. (**H**) Flow cytometry plot showing frequencies of HLA-G+ and CD49a+ cells from hTSC-TOrgs cultured in 21%, 3%, and 21%+ROX conditions following 7 days of differentiation (Left). Bar plots show the proportion of HLA-G+ cells and mature EVT (HLA-G+/ITGA1+) cells in 21%, 3%, and 21%+ROX on Day 7 of EVT differentiation (Right). Data plotted as mean values with standard deviation error. Statistical analyses were performed ANOVA and two-tailed Tukey post-test; differences significant at P<0.05. *= P<0.05,***= P<0.001.

In organoids cultured in 21% oxygen, spikey and elongated outgrowths were observed at endpoint, while outgrowth in 3% oxygen appeared rounded and less invasive (Fig. 5D). These findings are consistent with our prior observations (Fig. 3). Notably, the presence of ROX in 21% oxygen did not disrupt the spikey and elongated phenotype, suggesting that active HIF is not sufficient in blocking mature EVT differentiation (Fig. 5D). To examine this further, we measured gene markers of progenitor EVT (*ITGA2, TAGLN*), cCTB akin those in the proximal column (*NOTCH1*), and *HLA-G* (expressed in maturing cCTB and EVT) at days 0, 3, and 7 of EVT differentiation in vehicle and ROX treated organoids. On Day 0, organoids cultured in 3% O2 expressed higher levels of *ITGA2* and *NOTCH1* compared to ambient conditions (Fig. 5E), supporting our previous findings that low oxygen potentiates EVT progenitor development. At differentiation endpoint, progenitor EVT/cCTB markers were all greater in 3% oxygen than in 21% oxygen, further supporting our findings that low oxygen promotes the expansion of immature EVT (Fig. 5E). However, stabilization of HIF in ambient oxygen using ROX did not lead to differences in *ITGA2*, *TAGLN*, or *NOTCH1* levels at any differentiation timepoint when compared to 21% oxygen, though levels of *NOTCH1* at day 7 in ROX trended on being greater compared to untreated ambient conditions (Fig. 5E). Levels of *HLA-G* at differentiation endpoint are not altered by ROX (Fig. 5E). These findings were corroborated by quantifying the intensity of CD49b/*ITGA2* in trophoblasts cultured in ambient oxygen, 3% oxygen, or ambient oxygen/ROX combinations (Fig. 5F). While 3% oxygen led to a consistent increase in CD49b/ITGA2 levels (consistent with an expansion of progenitor EVT), HIF stabilization using ROX did not lead to CD49b/*ITGA2* induction (Fig. 5F), indicating that hypoxia contributes to the expansion of EVT progenitors independent of HIF. Together, these data suggest that HIF activity does not regulate the emergence of progenitor EVT (i.e., ITGA2^+^ cells), though HIF signaling may promote the formation of NOTCH1^+^ cCTB.

To better understand the relationship between the role of HIF activity and EVT maturation, we also examined levels of gene markers (*ITGA1*, *SERPINE2*, *NOTUM*) associated with mature EVT at Day 7 of EVT differentiation (Fig. 5G). Consistent with our single-cell transcriptomic data, 3% oxygen associated with markedly lower levels of all three mature EVT gene markers, indicating that persistent low oxygen significantly blocks the formation of mature EVT (Fig. 5G). While ROX-induced stabilization of HIF did not affect *NOTUM* levels compared to ambient oxygen, HIF stabilization did lead to a modest decrease in *ITGA1* and *SERPINE2* levels (Fig. 5G), providing evidence that HIF signaling may dampen EVT maturation. This interpretation is supported by the finding that HIF stabilization also dampened the formation of mature HLA-G+/ITGA1+ EVT compared to 21% oxygen; low oxygen (3%) led to the robust block of mature EVT (Fig. 5I, 5J).

## DISCUSSION

Prior to this work, our understanding of how low oxygen regulates EVT development was fragmented, largely because it was not possible to sequentially characterize the impact of oxygen on cells progressing along the EVT pathway at single-cell resolution. Using hTSC- and primary-derived trophoblast organoids combined with single-cell transcriptomics and trajectory analyses, we show that low oxygen (3–8%) selectively expands column-like cCTBs, stabilizing an immature EVT progenitor population, while restricting the formation of mature, invasive EVTs. In contrast, ambient oxygen (21%) promotes EVT maturation and villous SCT differentiation. Hypoxia-associated programs, including HIF1α/EPAS1 signaling, are activated during EVT differentiation independent of oxygen, yet HIF stabilization under 21% oxygen partially recapitulates the low-oxygen phenotypes, indicating that both HIF-dependent and HIF-independent mechanisms contribute to progenitor expansion and maturation restraint. Collectively, these findings provide a high-resolution framework for understanding how oxygen tension orchestrates the balance between progenitor maintenance and terminal differentiation along the EVT lineage, offering mechanistic insight into early placental development and potential dysregulation in pregnancy complications.

A paradigm exists that low oxygen promotes immature EVT formation while also leading to the restriction of more invasive mature EVT, as defined by the expression of HLA-G, ITGA5, and ITGA1, which together have been used to define the emergence of immature EVT (HLA-G, ITGA5) that reside in anchoring columns and invasive decidual-associated EVT (ITGA1)^26^. This has been suggested based on the piecemealing of studies spanning over twenty years of research that together create a somewhat contradictory interpretation of the role that low oxygen plays in controlling EVT development; that low oxygen tension both potentiates and impairs EVT differentiation. This current study, leveraging state-of-the-art trophoblast organoid platforms that accurately recapitulate the step-wise progression of EVT development with single-cell transcriptomic resolution verifies this untested paradigm. Our finding that low oxygen promotes initial EVT differentiation aligns closely with previous studies using primary 2D trophoblast cultures^16^ and placental explants^21,22^. Wakeland et al. showed that low oxygen promotes the transition of EGFR^+^ CTB into HLA-G^+^ proximal column-like EVT^16^. Additionally, our group as well as others have shown that exposure to low oxygen (1-3%) enhances column outgrowth in placental explant models^20–22^. Additionally, the observation that there is restriction of EVT maturation aligns with other work that show a low-oxygen dependent decrease in ITGA1+ expression^21,25^. Studies in the macaque also corroborate the importance of elevated oxygen tension in driving a mature and invasive trophoblast phenotype^41^.

The expansion of a cCTB-like population in low oxygen is interesting when considering the *in vivo* architecture of placental villi, and the local oxygen gradients present during development. Cells of the proximal and distal column of placental villi are likely experiencing a lower oxygen tension than the highly vascularized areas of the decidua, which contain the most invasive EVT. Additionally, the proximal column is thought to contain a EVT progenitor niche. Within this defined region the markers NOTCH1^10^, ITGA2^42^, SOX15^9^ and TAGLN^9^ identify these putative progenitors, with functional roles for establishing this progenitor population shown for NOTCH1. Although this area is broadly assumed to contain specifically EVT progenitors, our group as well as others have demonstrated that ITGA2^+^ cells may be bipotent, contributing to both EVT and SCT differentiation^9,42^. Progenitor maintenance under low oxygen conditions has been previously shown using the 2-D hTSC model^25^. The low oxygen environment present during early placental development may be important in maintaining or expanding this niche, while higher levels of oxygen, found deeper in the decidua, are needed for full EVT maturation. Interestingly, our results suggest that HIF-1 does not appear to be regulating this cCTB-like population, as stabilizing HIF signalling in 21% oxygen did not recapitulate the expanded progenitor EVT phenotype seen in low oxygen. Our studies demonstrate that HIF activity is not sufficient to drive EVT progenitor development; however, other studies have shown the necessity of HIF activity in controlling the development of HLA-G^+^ trophoblasts^16,17^. In our organoid system, it will be important to explore HIF necessity (i.e., depletion of HIF1a/2a or ARNT) in the control of discrete progenitor and column-like populations using single-cell transcriptomics. Additionally, it will be important to explore what low oxygen-dependent, HIF-1-independent genes and pathways may be regulating this population.

Our results suggest that HIF-1 is involved in the blunting of EVT maturation that is observed in low oxygen. Because HIF-1 largely acts as transcriptional activator not repressor, future work should focus on the which downstream targets HIF-1 may be acting through to cause this phenotype. Additionally, due to ROX stabilizing both HIF1α and EPAS1, the contribution of individual subunits on repressing maturation could not be explored. Delineating the contribution of individual subunits is also an important next step to better understand the role of HIF-1 in EVT differentiation.

### Limitations of the Study

scRNAseq analyses are limited by cell capture and relatively shallow read depth, so although we sought to mitigate these drawbacks by supporting our work with alternative approaches (bulk-RNAseq, microscopy, flow cytometry, etc.), these results should be interpreted with these caveats in mind. Cell capture limitations of the 10X Chromium platform related to the exclusion of large cells (> 40 um), like terminally differentiated EVT and multinucleated SCT, also necessitate overall tempering as the exclusion of such cell types excludes the modeling and assessment of trophoblast endpoint differentiation. Additionally, the pharmaceutical HIF stabilizer Roxadustat may induce unknown off-target effects that may complicate the interpretation of HIF-related findings. Lastly, our study is performed entirely *in vitro* applying rigid categories of oxygen tension that are not reflective of oxygen concentration gradients experienced within the placental bed.

## ACKNOWLEDGEMENTS

We thank members of the Beristain laboratory, specifically Maya Grayck and Elakkiya Prabaharan for their critical reading and feedback of this manuscript. We additionally thank the computational resources provided by the Advanced Research Computing Team, University of British Columbia. Thank you to the members of the Facility for Advanced Imaging and Microscopy at FMI, L. Gelman, L. Plantard, J. Eglinger, and T.-O. Buchholz for their support with imaging and quantification. We thank the functional genomics facility at FMI for performing bulk RNA sequencing experiments. We are grateful to Holly Anderson for the derivation of trophoblast organoids.

## AUTHOR CONTRIBUTIONS

Conceptualization: A.G.B., M.Y.T., G.L.M.; Methodology: G.L.M, G.G., I.C., A.H.K, H.H., K.K.; Formal analysis: bioinformatic – G.L.M., G.G., H.H., and benchwork – G.L.M., G.G., I.C., A.H.K, V.M.L, W.W, ; Resources: A.G.B., M.Y.T.; Writing: original draft – G.L.M., A.G.B.; review & editing – G.L.M., G.G., I.C., M.Y.T., A.G.B.; Supervision: A.G.B., M.Y.T., A.I.M.; project administration: A.G.B., M.Y.T.; Funding acquisition: A.G.B., M.Y.T, K.K.

## MATERIALS AND METHODS

### hTSC Trophoblast Organoid Methods

2-D hTSC Culture and Acclimation to Low Oxygen. Prior to organoid establishment, hTSC lines CT29 (RCB4937; male) and CT30 (RCB4938; female) were established as 2-D cultures in TS media. hTSC cultures were maintained on 10 μg/mL human placenta collagen IV (Sigma) in TS media containing DMEM/F12 (Gibco) supplemented with 0.1 mM 2-mercaptoethanol (Gibco), 0.2% Fetal Bovine Serum (Gibco), 0.5% Penicillin-Streptomycin (Gibco), 0.3% Bovine Serum Albumin (Sigma), 1% ITS-X supplement (Gibco), 1.5 μg/mL L-ascorbic acid, 50 ng/mL EGF (Thermofisher), 2 μM CHIR99021 (Sigma), 0.5 μM A83-01 (Sigma), 1 μM SB431542 (biogems), 0.8 mM 2-propylvaleric acid (Sigma), and 5 μM Y27632 (Sigma). To acclimate cells to 3% O_2_, cell lines were first grown in 21% O_2_ followed by passaging into 10%, 5%, and finally 3% O_2_ where they were maintained for 2-3 passages. All culture work below ambient oxygen was done in a Biospherix X3 physiological workstation.

#### Derivation and Culture of hTSC-Torgs

To establish hTSC-TOrgs from 2-D hTSC lines, cells were resuspended in undiluted growth factor-reduced Matrigel (Corning) and cell suspensions (40 μl) containing 5000 cells were seeded in each well of a 24-well plate. Plates were placed at 37°C for 2 min, followed by 15 minutes upside down to promote equal spreading of cells within each dome. Plates were turned back over and 0.5 mL of pre-warmed trophoblast organoid media containing advanced DMEM/F12 (Gibco), 10 mM HEPES (Gibco), 1 X B27 (Gibco), 1 X N2 (Gibco), 2 mM L-glutamine (Gibco), 100 μg/mL Primocin (Invivogen), 100 ng/mL R-spondin (Thermofisher), 1 μM A83-01 (Sigma), 100 ng/mL rhEGF (Thermofisher), 3 μM CHIR99021 (Sigma) and 2 μM Y-27632 (Sigma) was added to each well. This media is referred to as regenerative media.

#### hTSC-TOrg EVT Differentiation

hTSC-TOrgs were cultured in regenerative media until >50% colonies reached at least 100μm in diameter (∼6 days). Following this, organoids were maintained in EVT-differentiation media containing advanced DMEM/F12, 0.1 mM 2-mercaptoethanol (Gibco), 0.5% Penicillin-Streptomycin (Gibco), 0.3% Bovine Serum Albumin (Sigma), 1% ITS-X supplement (Gibco), 100 ng/mL NRG1 (Cell Signaling Technology), 7.5 μM A83-01 (Sigma), and 4% Knockout Serum Replacement (Gibco). Media was exchanged for EVT-differentiation media lacking NRG1 on day 5 of differentiation following visible trophoblast outgrowth. EVT differentiation was stopped on day 7 of culture for scRNAseq, qPCR, and flow cytometry analyses. For ROX experiments, 50μm of Roxadustat (Tocris) or DMSO (Sigma) vehicle control was added to EVT media (Figure 5B). Bright-field images of hTSC-TOrgs were taken on an AxioObserver inverted microscope (Zeiss) using a 5X objective.

#### Pimonidazole Assay

hTSC-TOrgs were established as previously described and cultured in regenerative media for a total of 10 days in 3%, 8%, or 21% O_2_. Two hours prior to collection, organoids were incubated with regenerative media containing 50 µM pimonidazole (Hypoxyprobe). Organoids were then incubated with 500 μl Cell Recovery Solution (Corning) for 40 min at 4°C. Free organoids were then transferred to tall-96-well polypropylene plates (2 ml volume per well, Axygen) using wide bore pipet tips. For cryosectioning, organoids were collected from wells using the automated liquid handler and dispensed as liquid-free piles on a stretched latex sheet (Rite Dent) such that when relaxed the area contracted from the size of a 96-well plate (7.5 x 13 cm) down to a microscope slide (25 x 30 mm). The resulting spheroid micro-arrays (SMAs) were then immediately frozen via a −30 °C aluminium block embedded in OCT sectioning medium (Fisher Scientific), as previously described^41,43^. Organoid cryosections (10 µm thick) were cut with a Cryostar HM560 (Microm International GmbH) cryostat, transferred to capillary-gap microscope slides (Fisher Scientific) and air dried and then immediately fixed in 10 % buffered formalin phosphate for 15 mins and transferred to PBS + 0.1 % Tween 20 (PBST). Slides were stained using paired-capillary gap slides, 120 µm gap, that drew up ∼ 200 µl solution. Prior to antibody staining slides were blocked in the PBST with 6 % normal goat serum for 15 min. Slides were stained for 1 h with 1:250 Rabbit-anti-pimo-FITC (Hypoxyprobe) along with the nuclear marker Hoechst 33342 (20 µg/ml, Sigma-Aldrich). Slides were then washed for 30 min and mounted with PBS prior to imaging. During all incubation steps, capillary gap slides were stacked and clamped in sets of up to 20 pairs and shaken at 300 rpm to reduce concentration dependent staining artefacts, using a reciprocating 2 cm stroke motion, along the shorter, 25 mm, axis of the slide surface. Slides were loaded onto a custom-built x-y fluorescence imaging system based on a Nikon Plan Fluorite Imaging 10X objective (Nikon) and a cooled PCO Edge 4.2 sCMOS camera (PCO) controlled using ImageJ software (ImageJ, https://imagej.nih.gov/ij/). Using this system, images of entire microscope slides were captured at a resolution of 0.65 µm/pixel, and reduced to 1.3 µm/pixel prior to analysis. Signal intensity was measured using ImageJ.

#### RNA isolation

Total RNA was prepared from hTSC-TOrgs using TRIzol Reagent (Invitrogen) followed by RNeasy Mini kit (Qiagen) clean-up following manufacturer’s protocol. RNA purity was confirmed using a NanoDrop Spectrophotometer (Thermo Scientific).

#### cDNA synthesis & qPCR

100 ng RNA was reverse transcribed using qScript cDNA SuperMix synthesis kit (Quantabio) and subjected to qPCR 2^−ΔΔCT^ analysis using PowerUP SYBR Green Master Mix (Applied Biosystems) on a StepOnePlus Real-Time PCR System (Applied Biosystems). All reactions were performed in triplicate and all values were normalized to endogenous 18S transcripts. Forward and reverse primers can be found in Table S4.

#### Immunoblotting

hTSC-TOrgs were incubated with 500μl Cell Recovery Solution (Corning) for 40 mins at 4°C to dissolve the Matrigel matrix. Organoids were then washed in ice-cold PBS. 200μL of 1X Lameili buffer (Thermo Scientific) was added directly to organoid pellets and incubated for 5 mins. Samples were immediately boiled and frozen to avoid protein degradation. For immunoblotting, cell protein lysates were resolved by SDS-PAGE and transferred to nitrocellulose membranes. Membranes were cut and probed using mouse polyclonal antibodies directed against HIF1A (1:500, MAB1536, R&D Systems) or EPAS1 (1:1000, D9E3, Cell Signalling Technology) and β-actin (1: 5000, Santa Cruz Biotechnology). Blots were imaged on a iBright Imaging System (Invitrogen).

#### Flow cytometry

Following a seven-day EVT differentiation (Figure 5B), hTSC-TOrg cultures were incubated with 500μl Cell Recovery Solution (Corning) for 40 mins at 4°C. Organoids were then washed in ice-cold PBS and dissociated into single cells using TrypLE Express (Gibco) supplemented with 5% DNAse I for 10 min at 37°C. Following this, cells were filtered through a 70μm Flowmi cell strainer. Once in suspension, cells were washed 2X with ice cold PBS and stained with Fixable Viability Dye 780 (1:1000; eBiosciences) for 20 min on ice. Cells were then washed in PBS and incubated with Fc Receptor Binding Antibody Inhibitor (1:20; eBiosciences) for 10 min at 4°C, followed by 30 min incubation with combinations of: APC anti-HLA-G (clone MEMG-9; 1:150; Abcam), PE anti-CD49b (clone P1E6-C5; 1:150; Biolegend), and BV421 anti-CD49a (clone SR84; 1:200; BD Biosciences). Cells were then washed in PBS and incubated with 2% PFA for 15 min at RT. Following fixing, cells were washed and resuspended in flow buffer (PBS supplemented with 2% Fetal Bovine Serum) and kept in the dark at 4°C until data acquisition. Data was acquired using a Becton Dickinson Biosciences FACSymphony A5 and data was analyzed using FlowJo v10.8.1.

#### scRNA-seq library construction

hTSC-TOrgs were established and EVT differentiated as previously described and three consecutive library constructions were carried out over seven days (Figure 1B). For “Day 0” samples, organoids were cultured for 6 days in regenerative media in 21% or 3% O_2_. For “Day 3” samples, organoids were further cultured in EVT media for 3 days in 3%, 8%, or 21% O_2_. For “Day 7” samples, organoids were further cultured in EVT media for an additional 4 days in 3%, 8%, or 21% O_2_. For all organoid collections, cultures were incubated with 500 μl Cell Recovery Solution (Corning) for 40 min at 4°C. Organoids were then washed in ice-cold PBS and dissociated into single cells using TrypLE Express (Gibco) supplemented with 5% DNAse I for 10 min at 37°C. Following this, cells were filtered through a 70μm Flowmi cell strainer. Single-cell preparations were stained with 7-Aminoactinomycin D (7-AAD) viability marker (Invitrogen) and sorted using a Becton Dickinson (BD) Biosciences FACS Aria. Prior to chip loading, cell preparations from CT29 and CT30 were pooled for each of the 8 samples. These cell suspensions were loaded onto the 10x Genomics Chromium Controller for GEM bead generation and single-cell barcoding. Libraries were prepared using the Chromium Next GEM Single Cell 3ʹ GEM, Library & Gel Bead Kit v3.1 (10x Genomics) following manufacturer’s protocol. Single-cell libraries were sequenced on an Illumina Nextseq2000 instrument at a sequencing depth of 40,000 reads/cell. Sequencing reads were aligned to *hg36* reference genome using cellranger v6.0.1. A total of 95,208 cells were captured for sequencing (21% n = 32,953, 3% n = 32,240, and 8% n= 30,015) at an average depth of ∼41,000 sequencing reads/cell across all 8 samples.

### SCT^OUT^ Trophoblast Organoid Methods

#### Derivation of Primary-Torgs

Tissue samples used for the generation of primary trophoblast organoids for this study were obtained from elective first-trimester pregnancy terminations from consenting patients at Addenbrooke’s Hospital, following the Declaration of Helsinki (2000) guidelines and under ethical approval from the Cambridge Local Research Ethics Committee (04/Q0108/23). This approval is now incorporated into the East of England–Cambridge Central Research Ethics Committee overarching ethics permission, granted to the Centre for Trophoblast Research biobank for the ‘Biology of the Human Uterus in Pregnancy and Disease Tissue Bank’ at the University of Cambridge (17/EE/0151). The established trophoblast organoid cultures for this study are authorized under a Material Transfer Agreements (MTA) between the University of Cambridge and the Friedrich Miescher Institute for Biomedical Research (Agreements No. G119629 and G108313).

#### Primary-TOrg (P-TOrg) Establishment

Primary-TOrgs were grown in Matrigel (Corning) droplets in regenerative trophoblast organoid medium. After 7 days P-TOrgs and Matrigel were mechanically disrupted using an electronic pipette for four cycles of 100 pipetting and then incubated with pre-warmed Accutase cell detachment solution for 5 mins at 37°C. Afterward, the suspension was manually dissociated by pipetting 100 times and filtered through a 40 µm cell strainer, the flow through was mixed with Matrigel. 25ul Matrigel domes are plated in 48well plate (Corning) and incubated at 37°C for 20min. After, 250ul of regenerative media was added per each well and medium was changed every 2-3 days. Primary-TOrgs were maintained in a humidified incubator at 37°C with 5% CO_2_ in either 21% O_2_ or 2-3% O_2_. A more extensive protocol is published^44^.

#### SCT^OUT^ Trophoblast Organoid generation

SCT^OUT^ primary-TOrgs were generated as previously described^28^. Briefly, after 5 days in culture, Matrigel was removed using Cell Recovery Solution (Corning). Trophoblast organoids were then plated in low attachment 96w plate (Corning) in suspension and cultured at 37°C for 3 days. For generating SCT^OUT^ hTSC-TOrgs the same protocol was followed but the organoids were switched to suspension culture after 4 days in Matrigel culture. Bright field images were taken using EVOS XL Core microscope (Invitrogen).

#### EVT Differentiation of SCT^OUT^ Trophoblast Organoids

SCT^OUT^ Primary- and hTSC-TOrgs were made to differentiate as previously described^28^. Briefly, SCT^OUT^ organoids at day 3 were filtered and organoids with size above 100 μm but smaller than 200 μm were plated on top of a 20 μL Matrigel coated 96w plate. Organoids were cultured in regenerative media for two days before switching to EVT media for 8 days [Advanced DMEM/F12 (Life Technologies), 0.1mM 2-mercaptoethanol (Gibco), 2 mM L-Glutamine (Life Technologies), 100ug/mL Primocin (Invivogen), 0.3% bovine serum albumin (Sigma), 1% ITS-X supplement (Gibco), and 4% knockout serum replacement (Thermo Fisher)]. Medium was changed every 2 days. Bright field images were taken using EVOS XL Core microscope (Invitrogen).

#### Immunofluorescence Imaging and Analyses

SCT^OUT^ Primary- and hTSC-TOrgs were fixed in 4% PFA (Thermo Fisher). Permeabilization was performed using 0.5% Triton for 45min. For SCT volume quantification, fixed and permeabilized TOs were plated on plasma treated and poly-L-lysine (Gibco) coated µ-Slide 8 Well Glass Bottom dishes (Ibidi) and left to stick for 1 hour before blocking. Blocking was performed using 3% Donkey Serum (Pan Biotech) in 1X PBS. Organoids were incubated with primary antibodies [SDC1 44F9 (Miltenyi Biotec), CDH1 24E10 (Cell Signalling Technology), HLA-G G233 (Novusbio)] in antibody buffer solution (3% Donkey Serum + 0.05% triton) overnight at 4°C. After four washes with 1X PBS, organoids were incubated with secondary antibodies in antibody buffer solution (3% Donkey Serum + 0.05% triton) overnight at 4°C or 3 hours at RT. After four washes with 1X PBS, imaging was performed confocal microscope Stellaris 5 (Leica). For SCT volume quantification, after immunostaining, organoids were cleared in a 50 % Ce3D solution [(N-methylacetamide (Sigma-Aldrich), Histodenz (Sigma-Aldrich), 1-Thioglycerol (Sigma-Aldrich) and Triton X-100] diluted in PBS overnight and imaged using a 2-Photon Stellaris 8 microscope with a 25X objective (Leica). To obtain the total volume of the SCT^OUT^ organoids, CDH1 and SDC1 signals were analyzed using ImageJ software, and SCT volume was determined by subtracting the total volume from the CDH1 volume. STB volume over the total organoid volume was then plotted as a ratio.

#### hCG ELISA

Supernatants and brightfield pictures of the wells were collected after three days in suspension, the conditioned media was stored at-80°C. Dilution factors were first determined on test samples before final experimentation. Manufacturers’ instructions were followed in the execution of the ELISAs. The calculated protein concentrations (μg/ml) were normalized for organoid confluency using the number of organoids per well.

#### SCT^OUT^ Primary-TOrg Passaging and Bulk RNA sequencing library preparation

For the bulk RNA sequencing experiment, P-TOrgs (n=2 donors) were dissociated every 9 to 10 days to obtain single cells. Between passaging steps, the organoid suspension was centrifuged for 6 minutes at 600×g. Organoids were collected and Matrigel was removed using Cell Recovery Solution (Corning) for 50 min on ice. They were then washed with cold PBS and mechanically broken by automatic (400 times) and manual (100 times) pipetting. Organoids were, first incubated with pre-warmed Accutase cell detachment solution (Gibco) for 5 mins at 37 °C and then, incubated with collagenase V (Sigma-Aldrich) diluted in 10% FBS/Advanced DMEM/F12 for 15 min at 37 °C. The digest was filtered through a 40 μm nylon mesh cell strainer to ensure single cell suspension, and the flow through was collected in falcon round bottom tubes (Corning). Cell pellets were diluted in regenerative organoid medium, and cells were counted using the NucleoCounter® NC-202™ Automated Cell Counter (Chemometec). 10,000 cells were added to 25 ul of Matrigel (Corning) and were grown at 37°C in a humidified atmosphere of 5% CO_2_ and 21% O_2_ or 2% O_2_. Samples for sequencing were collected after 4 or 6 rounds of single-cell dissociation and growth under the respective oxygen conditions. At the collection round, after 5 or 6 days in Matrigel, trophoblast organoids were switched to in suspension culture and maintained at 37°C in 21% or 2% O_2_ for 3 days.

#### RNA extraction

For bulk RNA sequencing, total RNA was extracted from SCT^OUT^ Primary-Orgs (n=2 donors). RNA was extracted from organoid pellets using the RNeasy Micro kit with on column DNase treatment (Qiagen), following manufacturer’s instructions. The RNA was resuspended in 14 μl RNase-free water. The purity and concentration of the RNA was determined using the UV-Vis Spectrophotometer NP80 (Implen).

#### Illumina stranded mRNA-seq library preparation and bulk RNA-sequencing

cDNA libraries were prepared using the Illumina Stranded mRNA-Seq (Illumina IDT DNA/RNA UDI) preparation kit according to the manufacturer’s instructions. Libraries were pooled and sequenced with the Illumina NovaSeq 6000 platform, producing paired end reads of 56 base pairs each (2x56bp). Demultiplexing and FASTQ file generation were performed using Illumina’s bcl2fastq2 software.

#### Bulk RNA-seq data analysis

Paired-end sequence reads were aligned to the human genome (GRCh38 primary assembly) using the“qAlign” function from the Bioconductor package QuasR version 1.48.1 0 with default parameters except for aligner = “Rhisat2”, splicedAlignment = “TRUE”, allowing only uniquely mapping reads. Raw gene counts were generated using the “qCount” function (QuasR) with GENCODE release 38 ‘basic gene annotation’ (“gencode.v38.annotation.gtf” as TxDb object) as query, with default parameters except useRead=first and orientation=opposite. The count table was modified by replacing the Ensembl gene ID with the gene symbol (if unique) or a combination of the gene symbol and the Ensembl ID (if the gene symbol was not unique) or keeping the Ensembl ID (for genes without a gene symbol) as the unique identifier.

### Single-cell RNA-seq data analysis

#### Data pre-processing & quality control

Cells were pre-processed using the Seurat R package (v5.0.1)^45^ ^45^, including the raw files from two publicly available datasets (available at: GEO174481 and E-MTAB-6701)^46,47^. To remove low quality cells, those cells containing <1250 and >7500 detected genes, <1000 and >80000 counts, and greater than 20% mitochondrial DNA content were excluded from analyses. After initial filtering, doublets were removed using scDblFinder (v1.16.0)^48^. Individual samples were scored based on expression of G_2_/M and S phase cell cycle gene markers, scaled, and normalized. Next, the “FindVariableFeatures” function was used with a “vst” selection method to prioritize highly variable genes in each sample for all downstream analyses. All samples were merged, split by an “Integration” variable (a combination of “Batch” and “Oxygen”), and integrated using the “SCTransform” v2 workflow^49^. After integration, data layers generated were combined into a single layer using the “JoinLayers” function.

#### Clustering & Cell Type identification

The top 50 principal components were used for initial dimensionality reduction and visualization using the Uniform Manifold Approximation and Projection (UMAP) algorithm. Because the publicly available in vivo datasets included non-trophoblast cells, the Seurat object was further filtered to exclude contaminating cells. Cells were selected if they expressed combinations of trophoblast lineage markers; KRT7, EGFR, TEAD4, TP63, TFAP2A, TFAP2C, GATA2, GATA3, HLA-G, ERVFRD-1 at a level greater than zero. Additionally, cells needed to not express the markers VIM, WARS, PTPRC, DCN, CD34, CD14, CD86, CD163, NKG7, KLRD1, HLA-DPA1, HLA-DPB1, HLA-DRA, HLA-DRB1, HLA-DRB5, and HLA-DQA1. All cells from the organoid samples were included regardless of gene expression as there was no risk of non-trophoblast cells in these datasets. Once all trophoblasts in the dataset were identified and subset, cells were re-clustered at a resolution of 0.18 using 30 principal components. Cell type annotation was completed by analyzing the differentially expressed genes identified using the “FindAllMarkers” function (Table S2) in combination with known trophoblast lineage markers. Cytotrophoblasts (CTB) were determined by expression of ELF5, PAGE4, and BCAM, cell column cytotrophoblasts (cCTB) by NOTCH1, TAGLN, and ITGA2, extravillous trophoblast (EVT) by HLA-G, ITGA1, and ADAM12, and SCTp by CYP19A1, ERVFRD-1, and CGB. Trophoblast projections were visualized in an integrated UMAP (Figure 1D).

#### Pseudotime Analysis

The Monocle3 package^50^ (v1.3.4) was used to explore the differentiation kinetics of EVT lineage cells in the organoid dataset (21%, 8%, and 3%). The EVT lineage subset (CTB 1, CTB 2, cCTB, EVT) was imported, clustered, and the principal graph was created using the “learn_graph” function. The root of the trajectory was determined programmatically and cells were ordered in pseudotime using the “order_cells” function. Calculated pseudotime values for each cell were visualized by projecting cells onto the UMAP embeddings, with colors representing pseudotime values for each cell. Pseudotime distributions along the predicted EVT trajectory were compared between different oxygen conditions (21%, 8%, 3%) using a density plots generated with ggplot2 (v3.3.4)^51^. Statistical significance was assessed using the Kruskal-Wallis Test from stats R package (v4.3.2).

#### Pathway analysis

The Single-Cell Pathway Analysis (v1.6.2) package^52^ was implemented to evaluate what signalling pathways were enriched in CTB 1 compared to EVT. The package was implemented using default settings with one modification. The “compare_seurat” function was adapted to increase the downsampling of the objects from 500 to 1000 cells per group. Gene signatures corresponding to Hallmark pathways were obtained from the msigdbr human datasets and formatted for the SCPA package using the “format_pathways” function. Comparisons were performed between CTB 1 and EVT cells for the 21%, 3%, and in vivo datasets. Pathways with an absolute fold change (FC) > 5 and adjusted p-value < 0.01 were identified as enriched in a particular cell type. Enriched pathways were visualized using ggplot2 (v3.3.4).

#### Differential gene expression

Differential gene expression analysis (DGE) was performed in Seurat using the “FindMarkers” function implemented on a MAST framework generated on the expression matrix data to compare CTB 1 and EVT. To include the whole cell transcriptome in DGE analysis, parameters were set to include: genes with a log_2_ fold-change difference between groups of “−INF”, genes with a minimum gene detection of “−INF” in all cells in each group, and genes with a minimum difference in the fraction of detected genes within each group of “−INF”. This list of genes was then filtered to include only genes contained within the Hallmark “Hypoxia” gene set^53^. The top 50 of those genes were plotted by absolute log_2_ fold-change in CTB 1 and EVT cells in 21% and 3% datasets (Figure 4D).

#### Statistical analysis

Data are reported as mean values with standard deviations. Analyses were carried out using GraphPad Prism (version 9.3.1). Data was first assessed for normality using a Shapiro-Wilk test. For single comparisons unpaired t-tests or non-parametric Mann-Whitney were performed. For multiple comparisons a two-way ANOVA followed by Tukey’s multiple comparison tests were performed. Differences were accepted as significant at P < 0.05.

## SUPPLEMENTAL FIGURE LEGENDS

**Supplemental Figure 1. Establishment of hTSC-TOrgs in discrete oxygen conditions and identification of trophoblast identity** (**A**) Representative immunofluorescent images showing pimonidazole and Hoescht 33342 signal in hTSC-TOrgs cultured in regenerative media for 10 days in 21%, 8%, or 3% O_2_. Black boxes depict inset of individual organoids from each group. Scale bars, 500µm and 200µm (inset). (**B**) Bar plot showing pimonidazole staining intensity of hTSC-TOrgs cultured in 21%, 8%, and 3% O_2_. Data plotted as mean values with standard deviation error. Statistical analyses between groups were performed using ANOVA and two-tailed Tukey post-test; differences significant at P<0.05. **= P<0.01 ****=P <0.0001. (**C**) Representative brightfield images of hTSC-TOrgs in 21%, 3%, and 8% O_2_ at Day 0 and Day 7 of EVT differentiation. Scale bars 100µm. (**D**) UMAP plots of trophoblasts from hTSC-TOrgs cultured in 21%, 8%, 3% oxygen or obtained from first trimester placenta single-cell datasets (*in vivo*). (**E**) Dot plot showing the average log expression (0-1) and percent expressed of CTB, cCTB, EVT, and SCT lineage markers within each trophoblast cluster in the in vivo and in vitro (21%, 8%, and 3% hTSC-TOrgs) datasets.

**Supplemental Figure 2. Evaluation of the impact of oxygen on primary-TOrg transcriptome** Dimensionality reduction PCA plot of bulk RNA-seq data derived from SCT^OUT^ primary-TOrgs cultured in either 2% or 21% oxygen. Samples are colour coded by condition; 2% (blue) and 21% (purple), and shapes represent donors; P-TOrg1(circle) and P-TOrg2 (square).

**Supplemental Figure 3. Low oxygen blunts TOrg EVT outgrowth (A)** Representative bright-field images of hTSC-TOrgs (CT29) cultured in 21% and 3% O_2_ on Day 0, Day 4, and Day 8 of EVT outgrowth assay. Scale bars, 200µm. (**B**) Representative bright-field images of Primary-TOrgs (pTOrg-1) cultured in 21% and 3% O_2_ on Day 0, Day 4, and Day 8 of EVT outgrowth assay. Scale bars, 200µm.

